# MLL3/MLL4 enzymatic activity shapes DNA replication timing

**DOI:** 10.1101/2023.12.07.569680

**Authors:** Deniz Gökbuget, Ryan M. Boileau, Kayla Lenshoek, Robert Blelloch

**Affiliations:** The Eli and Edythe Broad Center of Regeneration Medicine and Stem Cell Research, Center for Reproductive Sciences, University of California San Francisco, San Francisco, CA, USA; Department of Urology, University of California San Francisco, San Francisco, CA, USA; Helen Diller Family Comprehensive Cancer Center, University of California San Francisco, San Francisco, CA, USA; Department of Biomedical Engineering, Duke University, Durham, NC, USA

## Abstract

Mammalian genomes are replicated in a precise order during S phase, which is cell-type-specific^1–3^ and correlates with local transcriptional activity^2,4–8^, chromatin modifications^9^ and chromatin architecture^1,10,11,12^. However, the causal relationships between these features and the key regulators of DNA replication timing (RT) are largely unknown. Here, machine learning was applied to quantify chromatin features, including epigenetic marks, histone variants and chromatin architectural factors, best predicting local RT under steady-state and RT changes during early embryonic stem (ES) cell differentiation. About one-third of genome exhibited RT changes during the differentiation. Combined, chromatin features predicted steady-state RT and RT changes with high accuracy. Of these features, histone H3 lysine 4 monomethylation (H3K4me1) catalyzed by MLL3/4 (also known as KMT2C/D) emerged as a top predictor. Loss of *Mll3/4* (but not *Mll3 alone*) or their enzymatic activity resulted in erasure of genome-wide RT dynamics during ES cell differentiation. Sites that normally gain H3K4me1 in a MLL3/4-dependent fashion during the transition failed to transition towards earlier RT, often with transcriptional activation unaffected. Further analysis revealed a requirement for MLL3/4 in promoting DNA replication initiation zones through MCM2 recruitment, providing a direct link for its role in regulating RT. Our results uncover MLL3/4-dependent H3K4me1 as a functional regulator of RT and highlight a causal relationship between the epigenome and RT that is largely uncoupled from transcription. These findings uncover a previously unknown role for MLL3/4-dependent chromatin functions which is likely relevant to the numerous diseases associated with MLL3/4 mutations.

## INTRODUCTION

Chromatin exists at multiple structural levels which integrate to modulate gene activity. Recent evidence suggests that the temporal order by which DNA is replicated during S phase – known as DNA replication timing (RT) – is correlated with various chromatin features predictive of gene activity^13^. Measurements of RT across the genome reveals segments of relatively uniform RT (herein referred to as RT domains). On the level of chromatin structure, boundaries of frequently self-interacting domains defined by chromatin conformation capture assays – known as topologically associated domains (TADs) – align well with boundaries of RT domains^14,15^ and specific TAD subtypes were recently implicated in replication initiation^12^. Euchromatin-associated epigenetic histone modifications such as histone H3 acetylation are enriched in early replicating RT domains^9,16^, whereas heterochromatin associated histone H3K9 methylation is enriched in late replicating RT domains^1^. Similarly, RT domains harboring actively transcribed genes are generally replicated earlier than ones with inactive genes^2,4–8^. At the level of the whole nucleus, early replicating or active chromatin localizes to the center of the nucleus and late replicating or repressed chromatin localizes to the periphery of the nucleus^17–20^. While this evidence correlates RT with markers of gene activity, it remains unclear how these chromatin features causally and hierarchically relate to RT and how these relationships may change during cell state maintenance *vs* transition. Much of the evidence that correlates RT with selected chromatin features has been produced under steady-state conditions in unrelated cellular contexts with functional evidence limited to selected genomic loci^21,22^. These limitations complicate comparability and consolidation of the data into a general model. Here, we address these gaps by performing epigenomic profiling and using machine learning to unbiasedly evaluate 21 chromatin features for their ability to predict RT during steady-state and cellular differentiation. Functional validation of one of the most predictive marks, H3K4me1, uncovered a new critical role for histone monomethyltransferase MLL3/4 activity in the regulation of RT by locally promoting initiation of DNA replication.

## RESULTS

### Genome-wide changes in DNA replication timing during early ES cell differentiation

To understand how RT is regulated during cellular differentiation, we used an ES cell model following the transition from naïve to formative pluripotency, reflecting the physiologically relevant cell fate transition of the embryonic epiblast during peri-implantation period of mammalian development^23–27^. To measure genome-wide RT, we developed a fast and scalable EdU/biotin-labeling based approach (BioRepli-seq) based on relative enrichment of sequenced nascent DNA within 50kb genomic bins in early *vs* late S phase (Fig. 1a). Our results in naïve ES cells correlated well with previously published results using different methods, although BioRepli-seq showed a greater dynamic range across early to late RT (Extended Data Fig. 1a)^28^. Both naïve and formative pluripotency states showed clearly delineated early and late replicating domains that change in size and RT across megabases during differentiation (Fig. 1b). Principal component analysis attributes ∼80% of the variance in RT to the pluripotency state transition and confirms reproducibility of the data within cellular states (Fig. 1c). Heatmap and statistical analysis of genome-wide RT changes during pluripotency state transition reveals ∼30% of the genome changed RT (FDR < 0.05) with equal changes towards earlier and later RT (Fig. 1d,e). Together, these results reveal large changes in RT genome-wide during the relatively short developmental time window reflected by the naïve to formative pluripotency transition.

**Figure 1:**
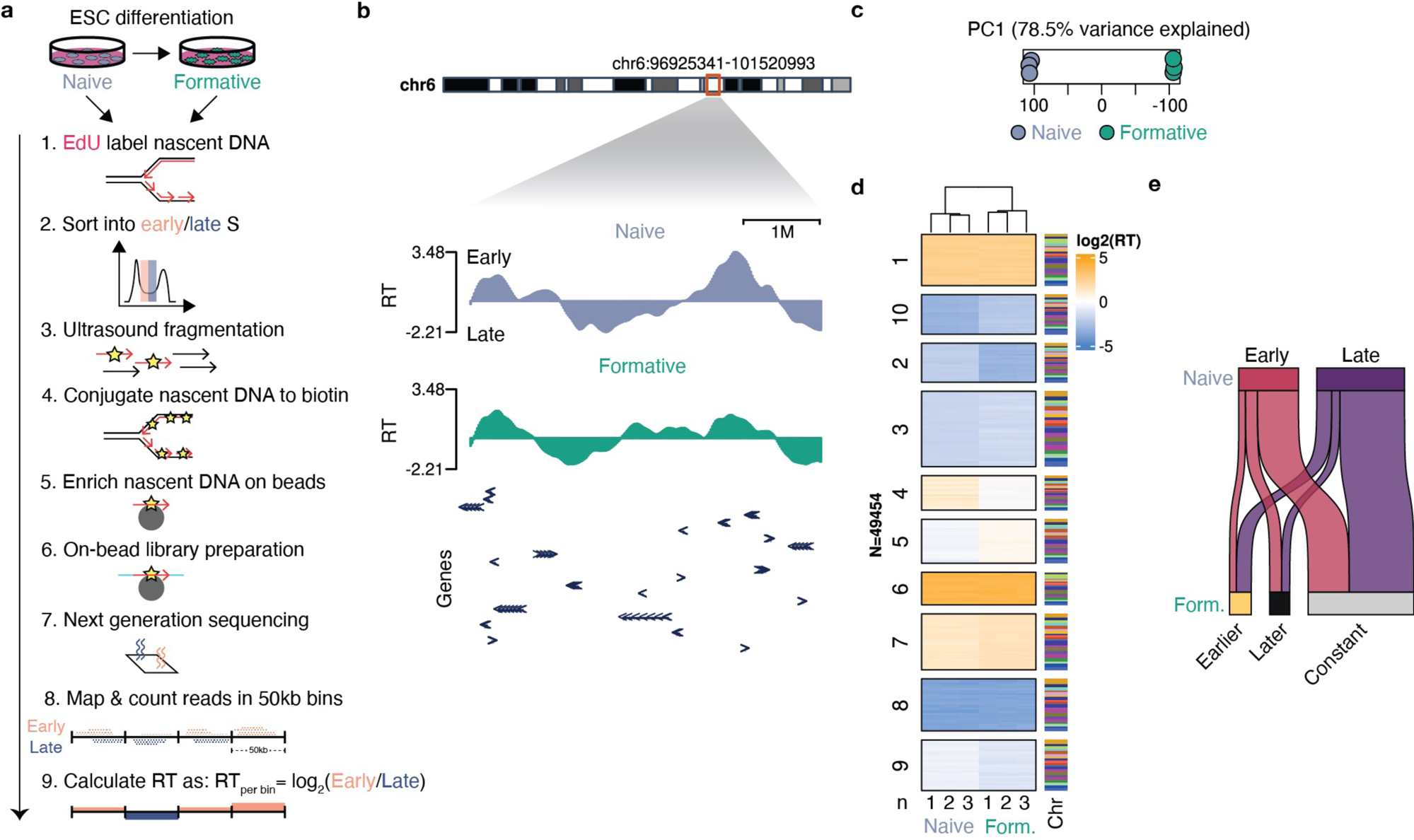
Genome wide changes in replication timing during pluripotency transition. **a**, Scheme of BioRepli-seq method and analysis. **b**, Genome track of log_2_ RT displaying early/late RT domains in naïve and formative state. **c**, Principal component analysis of genome-wide naïve (N) and formative (F) RT. Individual replicates derived from three independent cultures shown. **d**, Heatmap showing genome-wide RT per replicate of naïve to formative state. Associated chromosomes (Chr.) shown. Clustering performed by K-means. Individual replicates derived from three independent cultures shown. **e**, Sankey plot summarizing RT changes during pluripotency transition with absolute log_2_ fold change larger than 0.5.

### Chromatin state accurately predicts RT under steady-state and differentiation conditions

Next, we investigated the association between chromatin state and RT in steady-state naïve pluripotency, as well as how alterations in chromatin state are linked to changes in RT during the naïve to formative cell state transition. To do so, we measured the genome-wide signal of 21 chromatin features including histone modifications (H3K4me1-3, H3K27ac, H3K27me3, H3K36me1-2, H3K9ac, H3K14ac, H3K9me2-3, H4K8ac, H4K16ac, H4K20me1, H2BK5ac, H2AK119ub), histone variants (H3.3, H2Az, γH2AX), chromatin architectural factors (cohesin complex) and actively transcribing Pol II (phosphorylated at S2/5) using CUT&Tag under naïve steady-state conditions and following the naïve to formative cell state transition (Extended Data Fig. 1b,c). These data were processed and then used in machine learning models to determine which chromatin features were most predictive of naïve RT and changes thereof during differentiation (Fig. 2a, also see methods). Adjacent 50kb genomic bins were grouped into segments of similar naïve RT or delta (formative – naïve) RT (ΔRT) using circular binary segmentation^29^. Resulting segments for naïve RT and ΔRT had comparable median sizes of ∼1Mb, with 1894 naïve RT segments and 1077 ΔRT segments in total (Extended Data Fig. 2a,b). For each chromatin feature, we identified peaks using SEACR and summed the background-corrected peak signal within each RT segment. In the case of ΔRT segments, we calculated the difference between the summed signals for the formative and naïve steady states within each segment. The resulting summed chromatin feature signals within segments were then evaluated for their ability to predict steady-state naïve RT and ΔRT during differentiation using elastic net regression. The resulting models showed that chromatin features collectively were highly effective at predicting both steady-state naïve RT and RT changes during differentiation as evidenced by strong correlations of predicted *vs* observed RT (Fig. 2b,c). These data suggest a potential role for these chromatin features in driving RT both in steady-state and during cell state transition.

**Figure 2:**
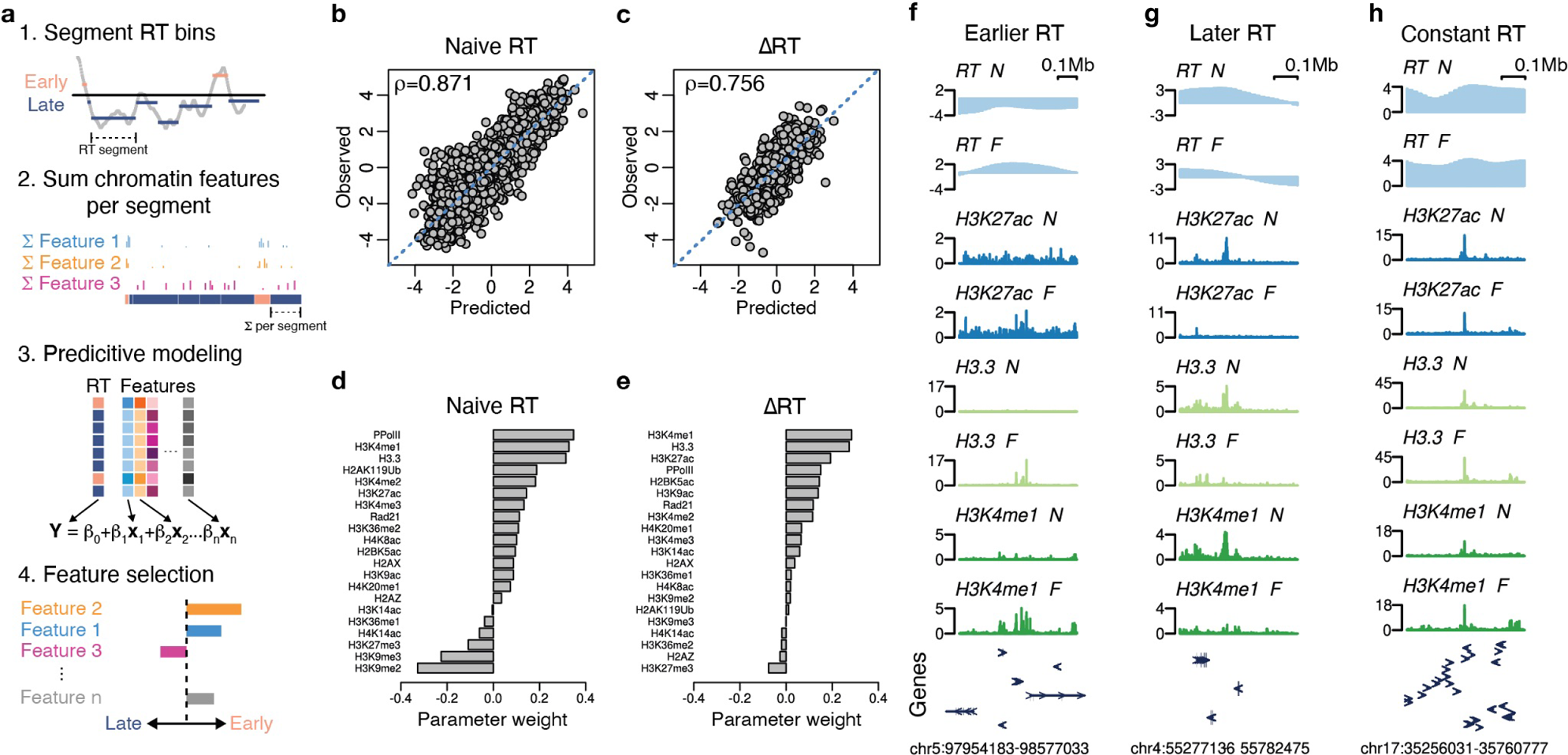
Chromatin state accurately predicts global RT with H3K4me1 emerging as top predictor. **a**, Workflow for RT segmentation, machine learning and feature selection. **b**,**c**, Scatter plots showing predicted *vs* observed naïve RT (b) and ΔRT (c) during transition based on fitted elastic net regression machine learning model. Spearman’s correlation coefficient (rho) shown. **d**,**e**, Parameter weight for each chromatin feature for naïve RT (d) and ΔRT (c) during transition based on fitted elastic net regression machine learning model. Larger magnitude of parameter weight implies more predictive strength of feature. **f**-**h**, Example genome tracks of earlier (f), later (g) and constant (h) RT domains and chromatin feature signal of top 3 predicting features in naïve (N) and formative (F) state. Exons depicted by arrowheads.

To better understand which of these chromatin features might be most influential on RT and changes in RT, we assessed the individual predictive strength of each feature by comparing their parameter weights in the generated regression models (Fig. 2d). Elastic net regression, a regularized machine learning model, shrinks the weights of parameters that offer redundant or no information to the model fit, facilitating the selection of uniquely predictive chromatin features. In the case of naïve RT, our analysis revealed strong positive parameter weights for phosphorylated Pol II, H3K4me1, and H3.3, along with strong negative weights for H3K9me2/3 and H3K27me3. For ΔRT during differentiation, change in H3K4me1, H3.3, and H3K27ac showed strong positive parameter weights, with only H3K27me3 showing a moderate negative parameter weight. The strong associations of H3K4me1, H3.3, H3K27ac, and RT were also readily apparent through visual examination of representative genome tracks (Fig. 2f-h).

To further validate these findings, we conducted linear regression for individual chromatin features and each pairwise combination. Correlation plots of genome-wide 50kb bins showed strong association of H3K4me1, H3.3, and phosphorylated Pol II signal intensity with early or, in case of differentiation, earlier RT (Fig. 2a,b). Linear regression analysis confirmed H3.3 and H3K4me1 as the top predictors of both naïve RT and changes in RT (see R^2^ values in Extended Data Fig. 2c,d). Specifically, H3.3 explained 56% and 32% of the variance in naïve RT and RT changes, respectively, while H3K4me1 explained 55% and 27%, respectively (Extended Data Fig. 2c,d). Furthermore, pairwise linear regression analysis of all marks demonstrated that the combination of H3.3 and H3K4me1 outperforms their individual predictive strength, validating that each mark provides separate information (Extended Data Fig. 2e,f). Similarly, both marks provide independent information from phosphorylated Pol II. In sum, these results provide quantitative evidence for H3K4me1 and H3.3 as strongest predictors of early/earlier RT in steady-state and during cell state transition. In contrast, while the repressive marks H3K9me2/3 and H3K27me3 are predictive of late RT in naïve ES cells, only H3K27me3 is moderately predictive of a shift to later RT during the transition.

### MLL3/4 activity is essential for genome-wide RT dynamics during ES cell differentiation

H3K4me1 was among the best overall predictors of early steady-state naïve RT and earlier RT changes during differentiation yet a causal role for this epigenetic histone modification or the responsible enzymes in the regulation of RT remains unexplored. H3K4me1 is commonly thought of as a marker of either primed or active enhancers that control the expression of their cognate genes^30^. The MLL3/4 (also known as KMT2C/D) enzymes are responsible for the H3K4 mono- and dimethylation at many of these enhancers^31,32^ with MLL4 being the predominantly active isoform in ES cells^33,34^. Surprisingly, the loss of MLL3/4 has little impact on the establishment of a formative transcriptional program during ES cell differentiation^34^. Furthermore, point mutation of the catalytic site of these enzymes resulting in the loss of their H3K4 methyltransferase activity has little to no impact on the transcriptional state in naïve steady-state or on transcriptomic changes during ES cell differentiation^34^. Hence, it remains unclear how these enzymes and their activity causally impact chromatin. We hypothesized that the regulation of RT may be one such role. To address this hypothesis, we performed BioRepli-seq in ES cells lacking MLL3 (MLL3^KO^), both MLL3 and MLL4 (MLL3/4^dKO^), or MLL3 and MLL4 catalytic activities (MLL3/4^dCD^) under both steady-state naïve conditions and following the naïve to formative pluripotency transition (Fig. 3a,b, Extended Data Fig. 3a-c). Principal component analysis confirmed reproducibility of the data across three replicates with clear separation between cell state and genotype. MLL3^KO^ overlapped with WT consistent with a primary role for MLL4 in these cells (Extended Data Fig. 3d,e).

**Figure 3:**
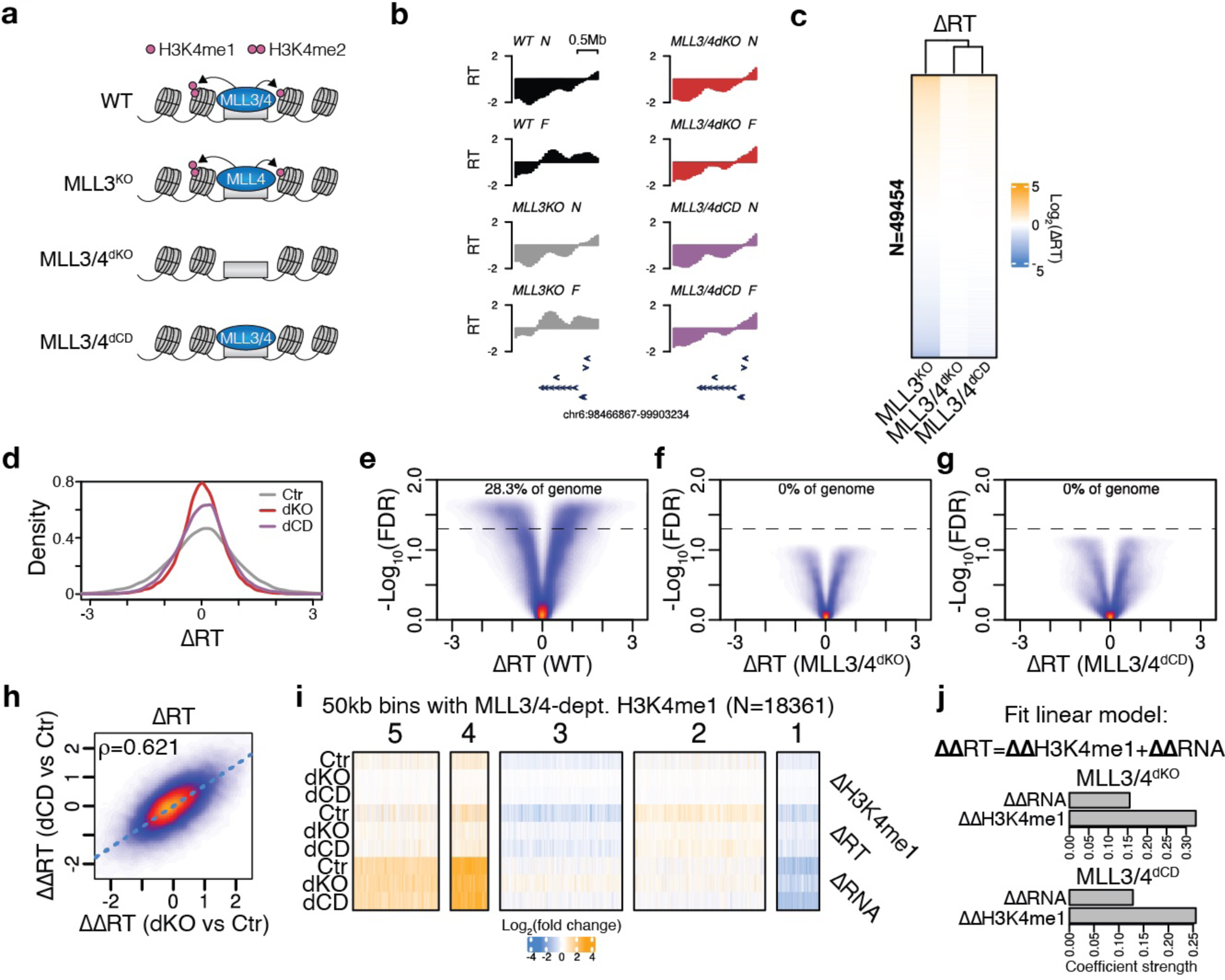
MLL3/4 activity shapes genome-wide RT dynamics during ES cell differentiation. **a**, Scheme of loss-of-function models. **b**, Example genome tracks of MLL3/4 activity-dependent RT domain showing average RT of WT, MLL3^KO^, MLL3/4^dKO^ and MLL3/4^dCD^ ES cells in naïve (N) and formative (F) state. **c**,**d**, Heatmap (c) and histogram (d) showing genome-wide ΔRT during pluripotency transition for control (MLL3^KO^), MLL3/4^dKO^ and MLL3/4^dCD^ ES cells. Heatmap sorted by control ΔRT. **e**-**g**, Volcano plots showing genome-wide ΔRT *vs* negative log_10_(FDR) during pluripotency transition for WT, MLL3/4^dKO^ and MLL3/4^dCD^ ES cells. Percentage of genome changing shown for FDR < 0.05. **h**, Correlation plot of ΔRT deficiencies in MLL3/4^dKO^ relative to controls vs MLL3/4^dCD^ relative to controls. Spearman’s correlation coefficient (rho) shown. **i**, Heatmap showing differential H3K4me1, RT and transcription during pluripotency transition for control (MLL3^KO^), MLL3/4^dKO^ and MLL3/4^dCD^ ES cells for all 50kb bins that display failure in achieving control RT changes in H3K4me1 in MLL3/4^dKO^ and MLL3/4^dCD^. Heatmap clustered by K-means. **j,** Coefficients of linear regression explaining difference in ΔRT between MLL3/4^dKO^ (top) or MLL3/4^dCD^ (bottom) relative control (MLL3^KO^) using respective differences in H3K4me1 and transcription.

Representative genome tracks showed that sites where RT normally shifts from late to early during the naïve to formative transition in WT and MLL3^KO^ cells failed to do so in MLL3/4^dKO^ and MLL3/4^dCD^ (Fig. 3b). To expand on this analysis, we compared genome-wide changes in RT across all 50kb bins between the naïve and formative states. Heatmap analysis of bins sorted by their RT change from earlier to later in MLL3^KO^ showed a dramatic reduction in RT dynamics in the two double mutant ES cells (Fig. 3c). This reduction was also evident in histograms showing reduced dynamic ranges of ΔRT in the double mutant ES cells (Fig. 3d). Indeed, while controls display significant changes in ΔRT during the transition at ∼30% of the genomic bins, MLL3/4^dKO^ and MLL3/4^dCD^ were almost devoid of significant changes (Fig. 2e-g). This dramatic loss of significant changes could not be explained by high variability among the mutant *vs* control replicates as principal component analysis showed strong alignment between the replicates in all states (Extended Data Fig. 3d,e). Furthermore, the deficiencies of MLL3/4^dKO^ and MLL3/4^dCD^ in ΔRT relative to controls during the naïve to formative transition was highly correlated (Fig. 3h). These data uncover a causal role for the MLL3/4 proteins and their enzymatic activity in regulating dynamic changes in RT during early ES cell differentiation.

### MLL3/4 shapes RT largely independent of transcription

The identification of MLL3/4 as a regulator of RT dynamics led us to investigate the extent to which these changes in RT may be linked to transcriptional regulation by MLL3/4. To address this, we compared changes in H3K4me1, RT, and RNA levels during the naïve to formative transition in MLL3/4^dKO^, MLL3/4^dCD^, and control ES cells, focusing on bins that showed MLL3/4-dependent H3K4me1. Clustering of the bins by K-means separated them into five groups, all of which showed a clear association between the loss of dynamic H3K4me1 and loss of RT dynamics in the double mutants. In contrast, changes in RNA levels showed little to no association with either H3K4me1 or RT at most sites. Consistent with this observation, linear regression analysis across all genomic bins showed that changes in H3K4me1 in MLL3/4^dKO^ or MLL3/4^dCD^ were much more predictive of changes in RT than changes in RNA levels (Fig. 3j).

We extended this analysis to steady-state conditions. MLL3/4^dKO^ and MLL3/4^dCD^ naïve ES cells showed large numbers of changes in RT relative to MLL3^KO^ controls (Extended Data Fig. 3f-h). Comparisons of changes in H3K4me1, RT, and RNA uncovered two clusters where the loss of H3K4me1 was associated with a shift to later RT (Extended Data Fig. 3i). Of these two clusters, only the smaller one representing many fewer genomic bins showed an association with changes in RNA levels. Linear regression across all genomic bins showed associations with both, but to a larger degree between RT and H3K4me1 than RT and transcription (Extended Data Fig. 3j). Together, these data underscore the genome-wide impact of MLL3/4 activity on RT and RT dynamics, which is largely uncoupled from transcription.

### MLL3/4 activity locally promotes DNA replication initiation

To investigate how MLL3/4 may facilitate earlier RT, we explored whether they directly promote initiation of DNA replication. Initially, we assessed subnuclear replication foci distribution using high-content microscopy at various stages of S phase. Replication foci were identified by briefly pulsing MLL3/4^dKO^ and control cells with the uridine analogue EdU, followed by fluorescent labeling and DNA co-staining. Utilizing an automated image processing pipeline, we identified nuclei and replication foci, categorizing them into quartiles based on sub-S phase DNA content (Extended Data Fig. 4a; see methods). Consistent with previous findings^35,36^, replication foci in control cells displayed a redistribution away from the nuclear center as S phase progressed (Extended Data Fig. 4c-d). In contrast, replication foci in MLL3/4^dKO^ cells did not exhibit such redistribution and remained significantly enriched closer to the nuclear center throughout S phase. These data indicate that MLL3/4 is required for the timely formation of more peripherally located replication foci that typically form in mid/late S phase.

To gain further mechanistic insights into how MLL3/4 may locally regulate DNA replication, we performed analysis of genomic regions that were specifically affected by MLL3/4 loss at the RT and H3K4me1 level. Genome tracks showed that these affected regions manifested as smaller, earlier-replicated peaks within megabase-scale late-replicated RT domains (Fig. 4a). The loss of MLL3/4 or their enzymatic activities led to the loss of these earlier RT peaks, resulting in large, consistently late-replicating domains. To gain insight into how MLL3/4 activity may promote earlier RT at these genomic sites, we next performed local metagene analysis of MLL3/4 ChIP-seq and H3K4me1 CUT&RUN data. In support of local MLL3/4 activity, MLL3/4 binding was enriched at the center of these peaks and coincided with MLL3/4-dependent H3K4me1 (Fig. 4b,c). We next evaluated whether the earlier RT within these peaks was driven by local MLL3/4-dependent DNA replication initiation zones (IZs). To do so, we mapped Okazaki fragment (OK)-seq and nucleoside analog incorporation loci (NAIL)-seq^38^ data derived from naïve ES cells to these MLL3/4-dependent RT regions. OK-seq involves strand-specific mapping of Okazaki fragments which allows detection of IZs based on Watson-Crick strand asymmetry. NAIL-seq, another method to detect IZs, is based on the sequential incorporation of two different nucleotide analogues and their subsequent signal subtraction resulting in signal enrichment at IZs. Analysis of OK-seq data demonstrated Okazaki fragment strand bias asymmetry at the center of MLL3/4-dependent RT peaks, indicating divergently moving DNA replication which is consistent with the presence of IZs (Fig. 4d). Further confirmation of the presence of IZs came from NAIL-seq and our EdU-seq data. Both datasets showed locally enriched signal which coincided with MLL3/4 binding and MLL3/4-dependent H3K4me1 (Fig. 4e,f). Strikingly, the loss of MLL3/4 proteins or their catalytic activity resulted in the loss of the EdU peaks (Fig. 4f). Together, these data support a local requirement for MLL3/4 activity in defining earlier RT by promoting IZs within large late-replicated RT domains.

**Figure 4:**
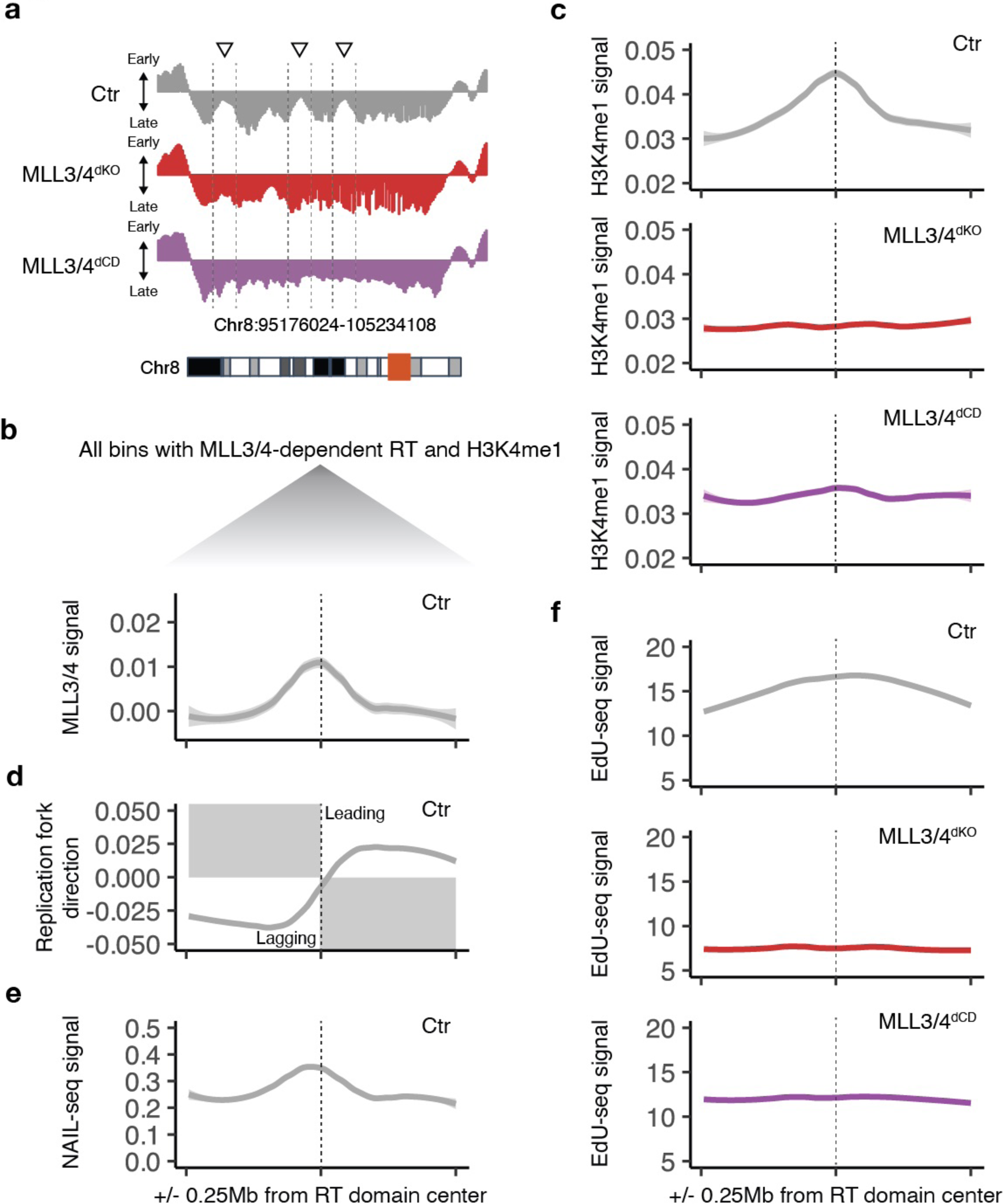
MLL3/4 activity locally promotes replication initiation. **a**, Example genome track of MLL3/4-dependent naïve RT domain for MLL3^KO^ (Ctr), MLL3/4^dKO^ and MLL3/4^dCD^ highlighting loss of early peaks withing larger later domain. **b**-**f**, Metagene analysis of WT MLL3/4 ChIP-seq (b), H3K4me1 CUT&RUN (for MLL3^KO^ (Ctr), MLL3/4^dKO^ and MLL3/4^dCD^) (c), WT OK-seq (d), WT NAIL-seq (e), and EdU-seq (for MLL3^KO^ (Ctr), MLL3/4^dKO^ and MLL3/4^dCD^) (f) centered at all MLL3/4-dependent naïve RT domains (log_2_ RT of MLL3/4^dKO^/Ctr > 2) that also show loss of H3K4me1 (log_2_ fold change of MLL3/4^dKO^/Ctr < −1).

### MLL3/4 promote MCM2 recruitment

To further investigate the potential mechanistic role of MLL3/4 in driving IZs, we categorized all H3K4me1 peaks in naïve ES cells into MLL3/4-dependent and -independent peaks, followed by metagene analysis (Fig. 5a). As anticipated, sites of MLL3/4-dependent peaks, unlike independent ones, exhibited a strong coinciding MLL3/4 binding signal (Fig. 5b). Analysis of OK-seq revealed a more prominent strand bias asymmetry at sites of MLL3/4-dependent *vs* -independent H3K4me1 peaks indicative of more pronounced firing of IZs (Fig. 5c). Similarly, NAIL-seq data displayed a stronger peak at sites of MLL3/4-dependent *vs* -independent H3K4me1 peaks (Fig. 5d). These data demonstrate that MLL3/4-dependent H3K4me1 peaks are enriched within IZs at a genome-wide level.

**Figure 5:**
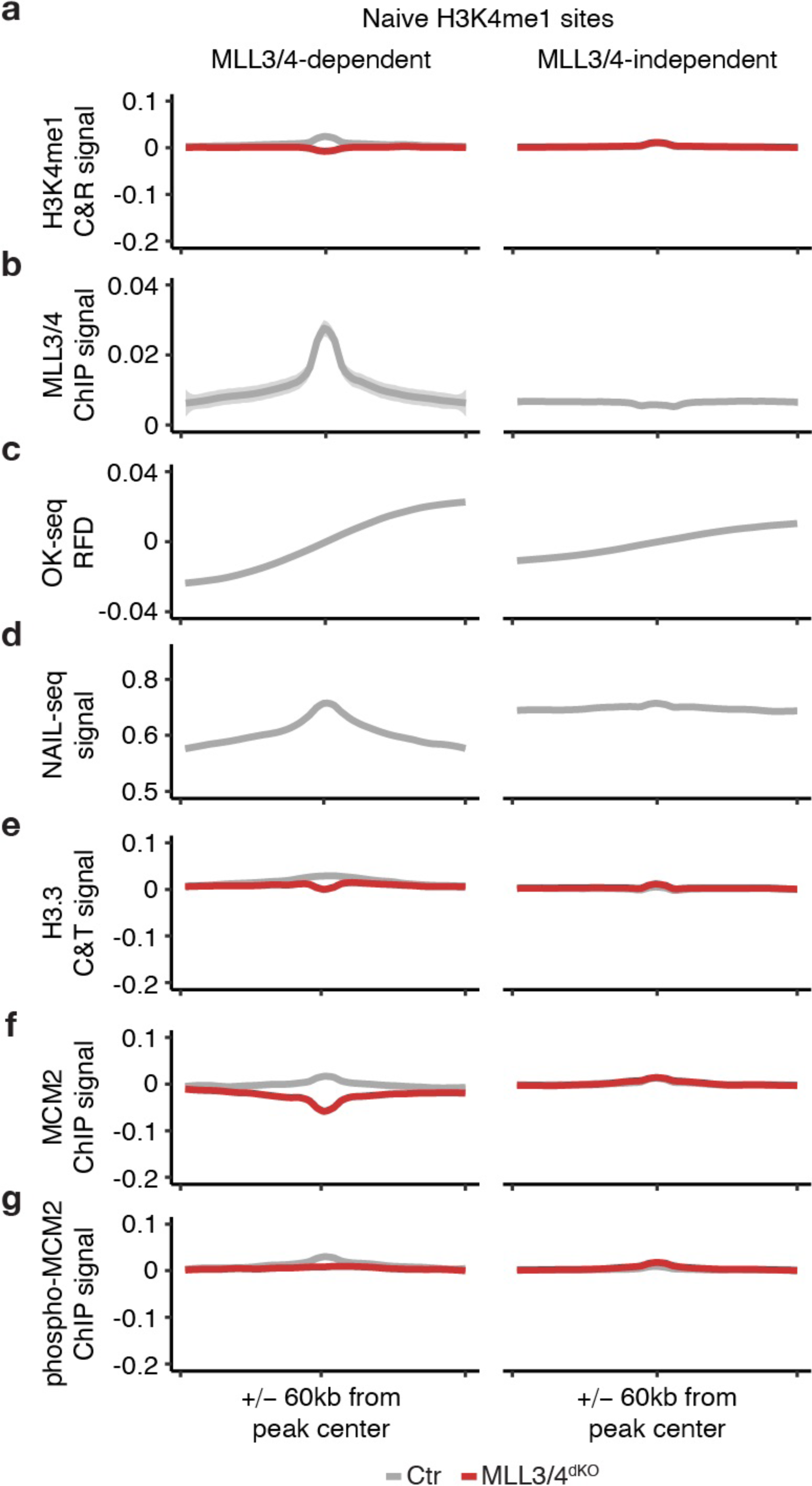
MLL3/4 promotes replication origin licensing and H3.3 recruitment. **a**-**g**, Metagene analysis of H3K4me1 CUT&RUN (for MLL3^KO^ (Ctr) and MLL3/4^dKO^) (a), WT MLL3/4 ChIP-seq (b), WT OK-seq (c), WT NAIL-seq (d), H3.3 CUT&Tag (for MLL3^KO^ (Ctr) and MLL3/4^dKO^) (e), MCM2 ChIP-seq (for MLL3^KO^ (Ctr) and MLL3/4^dKO^) (f), and phosphorylated MCM2 ChIP-seq (for MLL3^KO^ (Ctr) and MLL3/4^dKO^) (g) centered at all MLL3/4-dependent and -independent H3K4me1 peaks. H3K4me1, H3.3, MCM2 and phosphorylated MCM2 signal was normalized to local background to allow comparison of peak intensities across treatments.

Histone variant H3.3, like MLL3/4-dependent H3K4me1, marks enhancers^30,39,40^. Recent evidence indicates that H3.3 also can play a role in promoting earlier replicating IZs in the cancer cell line HEK293 under steady-state conditions^41^. Notably, in our predictive modeling, H3.3 emerged as a top predictor for both steady-state naïve RT and ΔRT during the transition (Fig. 2d-f). To explore whether H3.3 acts downstream of MLL3/4, we conducted CUT&Tag for H3.3 in MLL3/4^dKO^ and control ES cells. In controls, H3.3 signal was slightly more enriched at sites of MLL3/4-dependent *vs* -independent H3K4me1 peaks (Fig. 5e). Loss of MLL3/4 led to a specific depletion of H3.3 at dependent sites. These results support an upstream role for MLL3/4 on local H3.3 recruitment and suggest that MLL3/4 could be acting through H3.3 to promote earlier IZs.

Finally, we sought to determine how MLL3/4 may promote IZs by evaluating potential roles in the establishment of replication origins. Eukaryotic replication origins undergo a two-step activation process: 1) the formation of a pre-replication complex (origin licensing) involving the recruitment of the MCM proteins and 2) the formation of the pre-initiation complex (origin firing) requiring phosphorylation of the recruited MCM proteins^42^. To determine whether MLL3/4 contributes to origin licensing or activation, we conducted ChIP-seq for MCM2 and phosphorylated MCM2 (S40P) in MLL3/4^dKO^ and controls ES cells. Both MCM2 and phosphorylated MCM2 were enriched *vs* IgG control at the IZs identified by OK-seq (Extended Data Fig. 5). Metagene analysis at sites of MLL3/4-dependent and -independent H3K4me1 peaks revealed an enrichment for both MCM2 and phosphorylated MCM2 at both sets of sites (Fig. 5f,g). However, loss of MLL3/4 resulted in a specific loss of this enrichment at MLL3/4-dependent but not independent sites. Intriguingly, H3K4me1, H3.3, and MCM2 peaks were not only lost at dependent sites but also appeared locally depleted, suggesting that MLL3/4 loss may render local chromatin less accessible in addition to the failed deposition of these marks. These data uncover a role for MLL3/4 in the establishment of IZs through the recruitment of MCM proteins, contributing to the licensing of replication origins.

## DISCUSSION

Our results present a previously unknown role for MLL3/4 histone monomethyltransferase activity in the regulation of RT. In the absence of the MLL3/4 proteins or their enzymatic activity, genomic regions that are typically associated with dynamic H3K4me1 during the pluripotency transition show genome-wide coordinated loss of H3K4me1 and RT dynamics. Additionally, a subset of these domains exhibits coincident loss of H3K4me1 and delay of RT under naïve steady-state conditions. These affected regions constitute earlier replicated domains located within larger late replicating domains and enclose MLL3/4-driven IZs. MLL3/4 appears to achieve this function by facilitating recruitment of MCM proteins and thereby contributing to replication origin licensing (Fig. 6).

**Figure 6:**
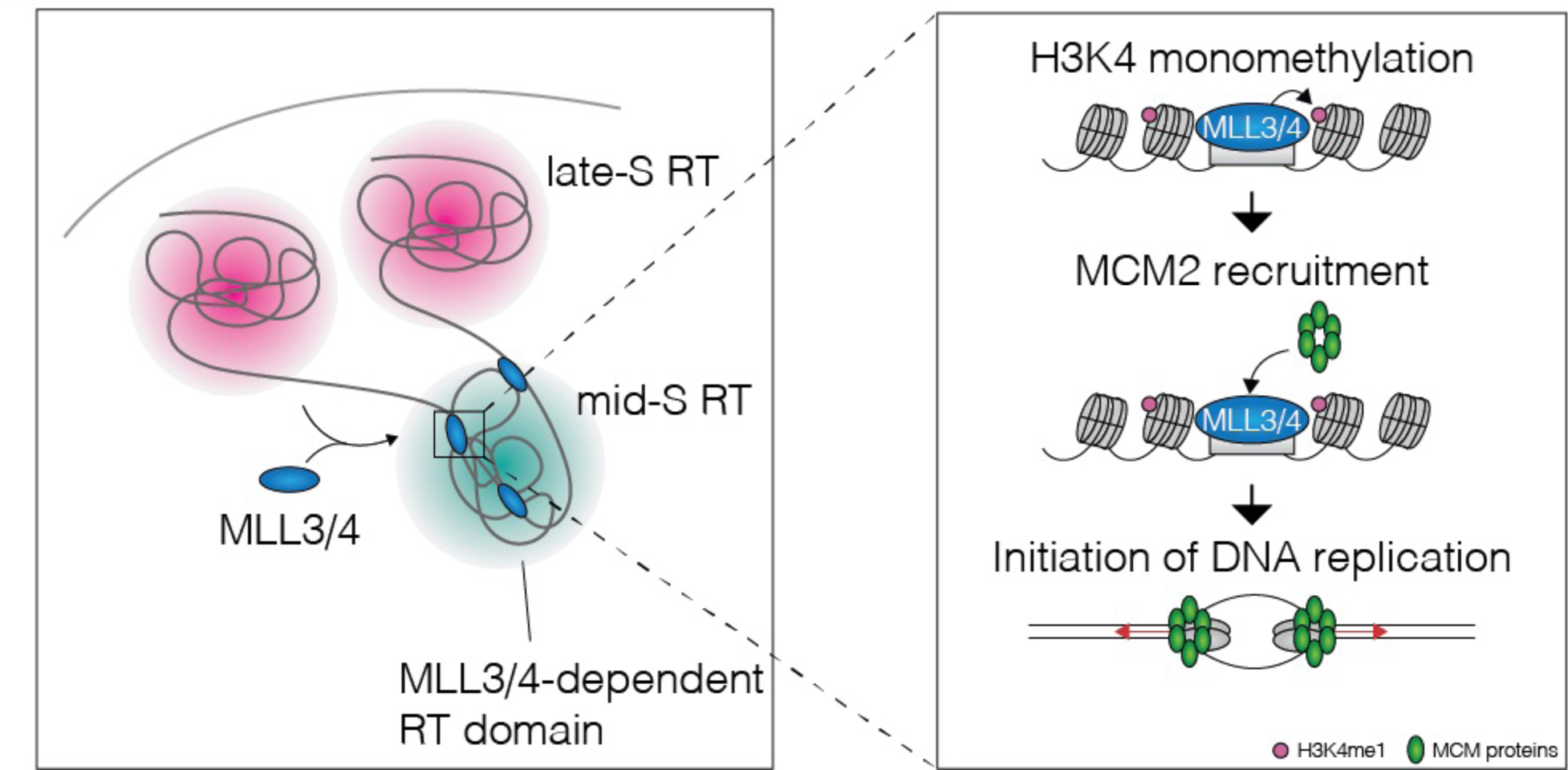
Model of MLL3/4 functions.

MLL3/4-driven H3K4me1 was one of the best predictors of RT in our analyses. Another strongly predictive mark, H3.3, was recently demonstrated to play a crucial role in the timing of early replication domains under steady state in the transformed cell line HEK293^41^. In this context, we present evidence indicating that H3.3 is downstream of MLL3/4, implying it may serve as an intermediate step between MLL3/4 and the establishment of RT. Notably, several chromatin features that demonstrated predictive power for RT under steady-state conditions exhibited reduced predictive strength during the transition from naïve to formative cell states. This observation suggests a potential distinction between bystander and driver-associated features. Overall, we find that chromatin state was highly predictive for RT, and the coordinated genome-wide loss of H3K4 mono/dimethylation and RT dynamics upon MLL3/4 inactivation suggests a functional coupling between the two processes.

MLL proteins are canonically thought to be directly involved in transcriptional regulation through deposition of active chromatin marks H3K4me3 at promoters by MLL1/2 and H3K4me1 at enhancers by MLL3/4^30,43^. While the proteins and their homologues are essential for embryogenesis and cell growth across multiple species^44–49^, the developmental relevance of their enzymatic activity seems to be more species specific. In flies, loss of H3K4 monomethyltransferase activity by mutation of the enzymatic domain of Trr is compatible with development and results in minimal gene expression changes^50^. However, mutant mice expressing enzymatically inactive MLL3 and MLL4 die during early development at embryonic day 6.5^47^. Despite the importance of MLL3/4 proteins and their enzymatic activity in mammalian development, the molecular basis for their requirement has remained elusive. While a role in transcription may in part explain their requirement, recent results suggest it is unlikely to explain the developmental phenotype. For example, MLL3/4 proteins were shown to drive all dynamic enhancer H3K4me1 during the naïve to formative pluripotency transition and their loss changes cell morphology and slows proliferation. Yet, expression of nearby genes was little to not impacted^51^. This result strongly suggests additional molecular roles for MLL3/4 proteins. Our unexpected discovery of a critical role for MLL3/4 methyltransferase activity in the regulation of RT might at least in part explain the presence of cellular and developmental phenotypes despite a paucity of direct transcriptional changes. Also, where there are transcriptional impacts, MLL3/4-driven changes in RT may be the culprit by rendering chromatin compartments permissive or repressive to gene activity.

RT is a consequence of timely activation of replication origins during S phase. Firing replication origins are enriched at promoters and enhancers^52^. We demonstrated that enhancer mark H3K4me1 is highly predictive of RT and that loss of its depositing enzymes MLL3/4, or their enzymatic activities impairs firing of replication initiation zones at MLL3/4-dependent H3K4me1 sites. MLL3/4 facilitates recruitment of both MCM2 and phosphorylated MCM2, which is consistent a role in origin licensing. A subset of earlier firing replication origins was recently shown to depend on cohesin-mediated 3D chromatin interactions^12^. Intriguingly, MLL3/4-dependent H3K4me1 can facilitate cohesin recruitment^53^ raising the possibility that MLL3/4 activity may exert its function to promote origin licensing through facilitating 3D chromatin interactions. Temporally, such a mechanism would be feasible as both H3K4 methylation^54^, and 3D chromatin interactions^55,56^ are rapidly reestablished after mitosis in early G1 simultaneously to when RT has been shown to be determined^57^.

*MLL*3 and *MLL*4 are both among the 10 most mutated genes in cancer (20% of all cancers)^58^. Our results present MLL3/4 as one of few known regulators of RT expanding its functional spectrum beyond transcriptional regulation. Whether changes in RT could in part explain the close association of mutations of MLL3/4 with cellular transformation will be important to investigate. Our results also uncover the close association of the epigenome with RT that can at least in part be uncoupled from transcription and more directly linked to origin firing. As such, our findings provide an important addition to the framework of how chromatin shapes cell identity.

## ACKNOWLEDGEMENTS

We thank Dr. Dan Wagner and Hannah Driks for critical reading of the manuscript draft. Technical support by the UCSF Parnassus Flow Cytometry Core and the UCSF Center for Advanced Technology is greatly appreciated. This publication includes data generated at the UC San Diego IGM Genomics Center utilizing an Illumina NovaSeq 6000 that was purchased with funding from a National Institutes of Health SIG grant (#S10 OD026929). R.B. would like to dedicate this paper Jean-Michel Vos, an excellent scientist and mentor who left this world far too early. This research was supported by NIGMS funding to RHB (R01GM122439 and R01GM125089).

## AUTHOR CONTRIBUTIONS

Conceptualization: DG, RB; methodology: DG; validation: DG; formal analysis: DG; investigation: DG, RMB, KL; resources: RB; data curation: DG; writing - original draft: DG; writing - review & editing: DG, RB; visualization: DG; supervision: DG, RB; project administration: RB; funding acquisition: RB.

## EXTENDED DATA FIGURES

**Extended Data Figure 1:**
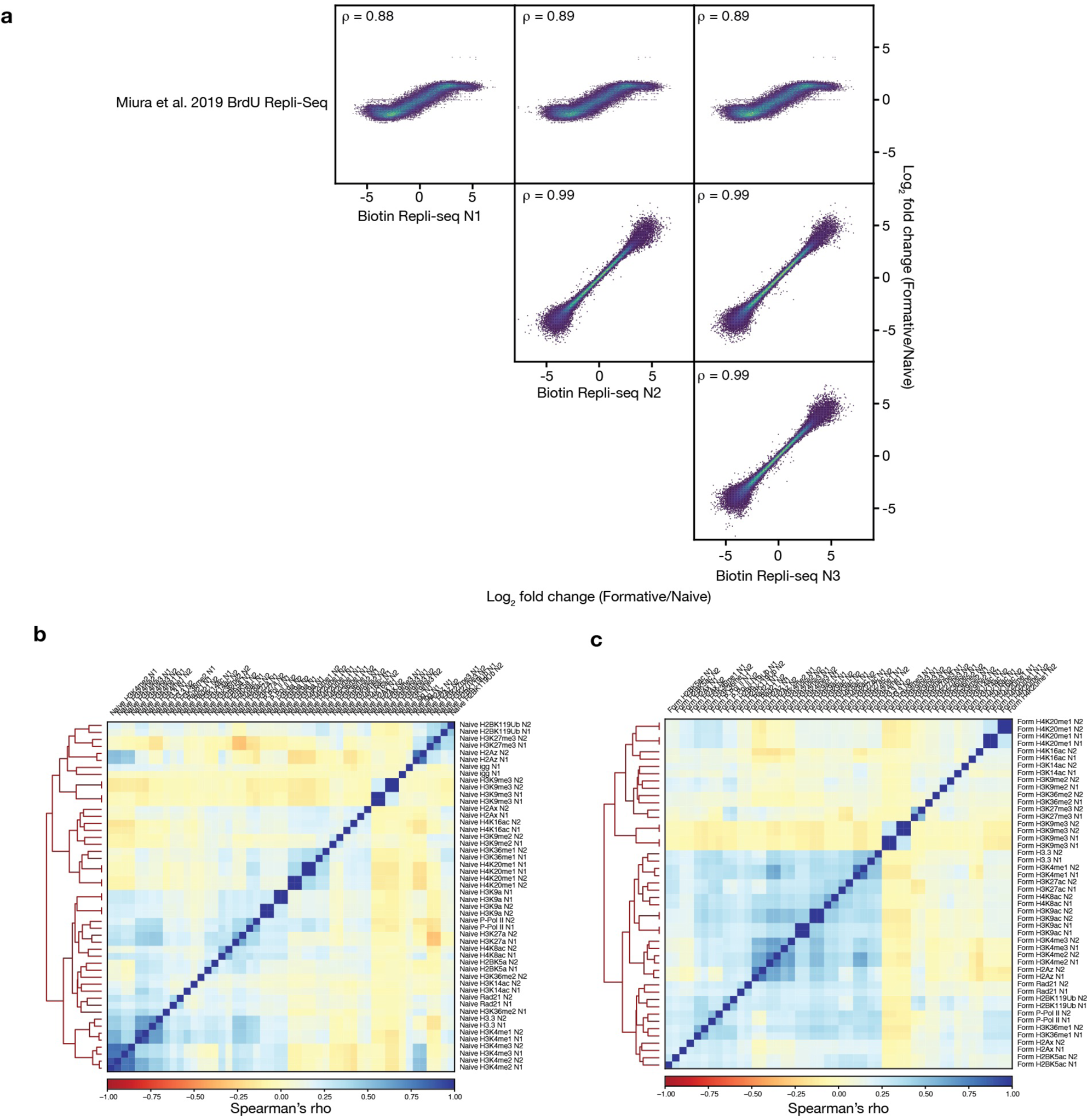
Quality control of BioRepli-seq and CUT&Tag. **a**, Correlation plots of BioRepli-seq replicates (derived from independent cultures) against each other and against previous BrdU Repli-seq method^28^. **b**, Correlation matrix of chromatin feature CUT&Tag data with individual replicates (derived from independent cultures).

**Extended Data Figure 2:**
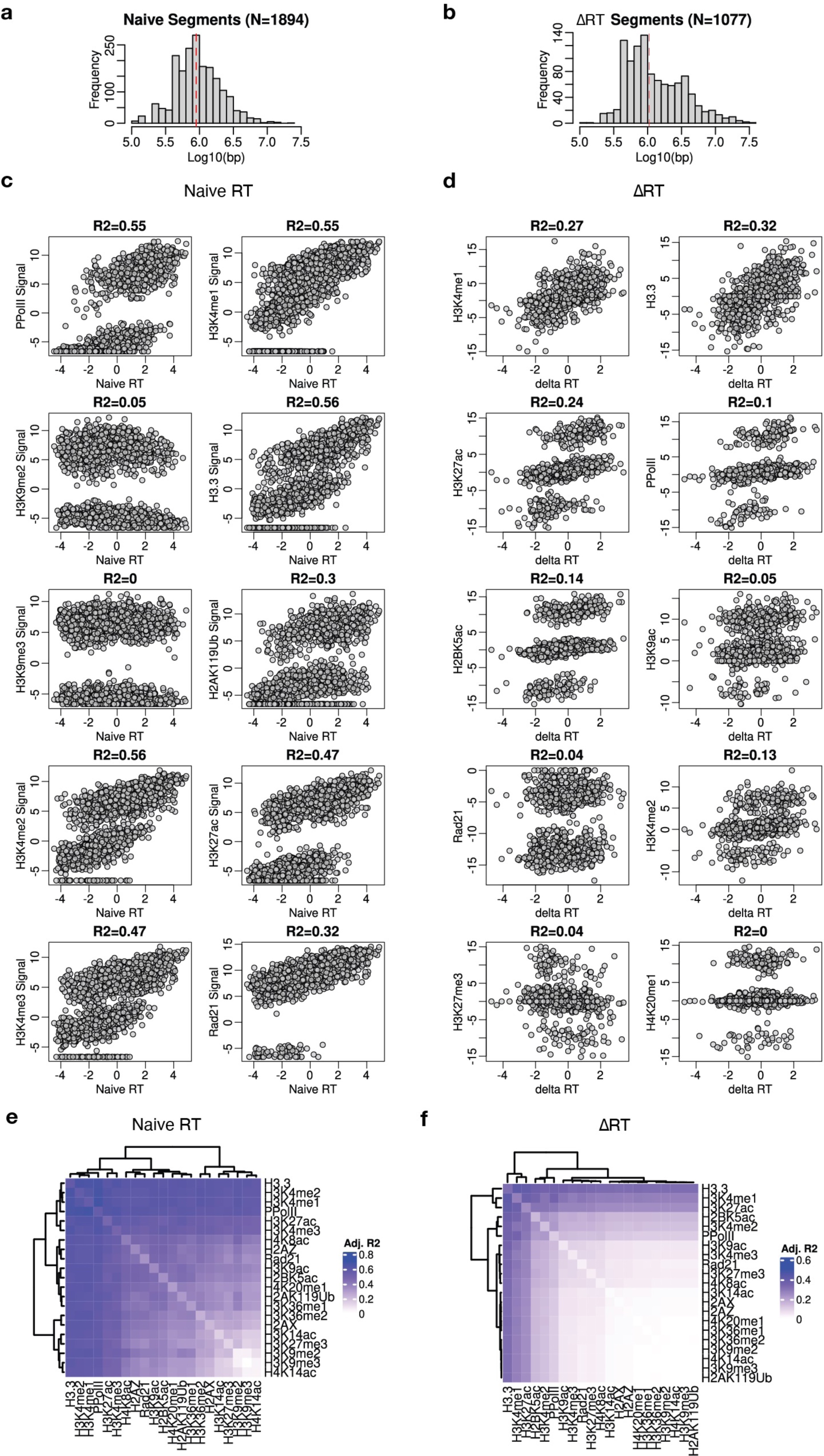
Associations of chromatin feature signal and RT. **a**,**b**, Histograms showing naïve RT (a) and ΔRT (b) segment sizes. Dashed line represents median. **c**,**d**, Scatter plots of naïve or differential chromatin feature signal *vs* naïve RT or ΔRT, respectively, for top 10 most predictive chromatin features based on elastic net regression model parameter weights (see Fig. 2d,e). Adjusted R^2^ derived from linear regression shown. **e**,**f**, Heatmap of adjusted R^2^ derived from pairwise or individual (diagonal) linear regression of naïve or ΔRT using naïve or differential chromatin feature signal, respectively.

**Extended Data Figure 3:**
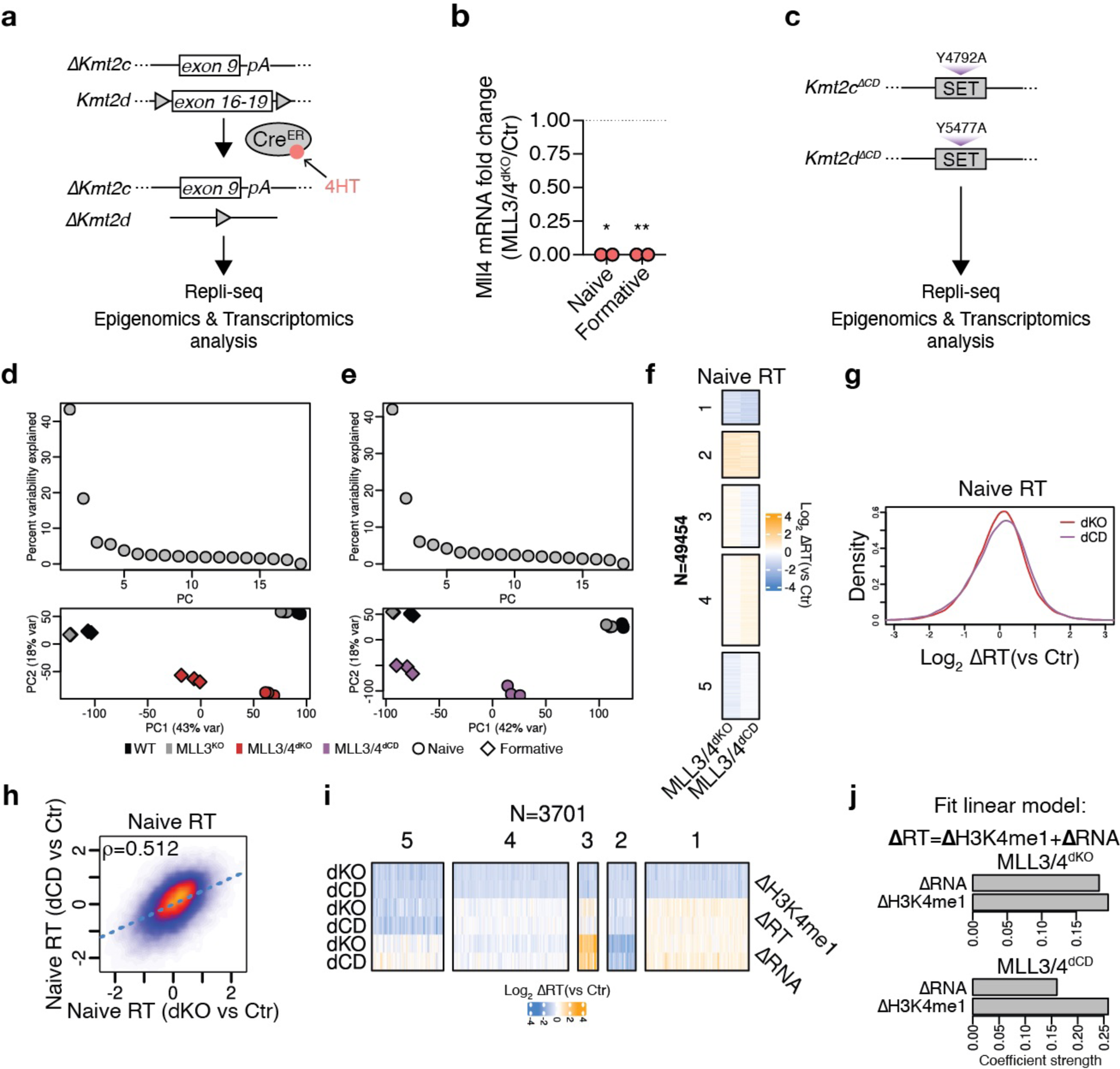
MLL3/4 activity shapes naïve RT. **a**, Scheme of genetics of MLL3/4^dKO^ ES cells. **b**, Relative Mll4 mRNA levels in MLL3/4^dKO^ *vs* Ctr at naïve and formative states measured by qRT-PCR. Gapdh mRNA was used as a reference. **c**, Scheme of genetics of MLL3/4^dCD^ ES cells. **d,e**, Principal component analysis of naïve (d) and formative RT (e) of WT, MLL3^KO^ and MLL3/4^dKO^. Individual replicates derived from independent cultures shown. Variance explained per PC (top) and PC1 *vs* PC2 shown (bottom). **f,g**, Heatmap (f) and histogram (g) showing genome-wide RT changes for MLL3/4^dKO^ and MLL3/4^dCD^ relative to control (MLL3^KO^) in naïve ES cells. Heatmap clustered by K-means. Individual replicates derived from three independent cultures shown. **h**, Correlation plot comparing RT changes in MLL3/4^dKO^ *vs* MLL3/4^dCD^ relative to control in naïve ES cells. Spearman’s correlation coefficient (rho) shown. **i**, Heatmap showing changes for H3K4me1, RT and transcription in MLL3/4^dKO^ and MLL3/4^dCD^ relative to control ES cells for in naïve ES cells for all 50kb RT bins that display reduction in H3K4me1 in MLL3/4^dKO^ and MLL3/4^dCD^ relative to control ES cells. Heatmap clustered by K-means. **j,** Coefficients of linear regression explaining changes in RT in MLL3/4^dKO^ (top) or MLL3/4^dCD^ (bottom) relative to control in naïve ES cells using respective changes in H3K4me1 and transcription.

**Extended Data Figure 4:**
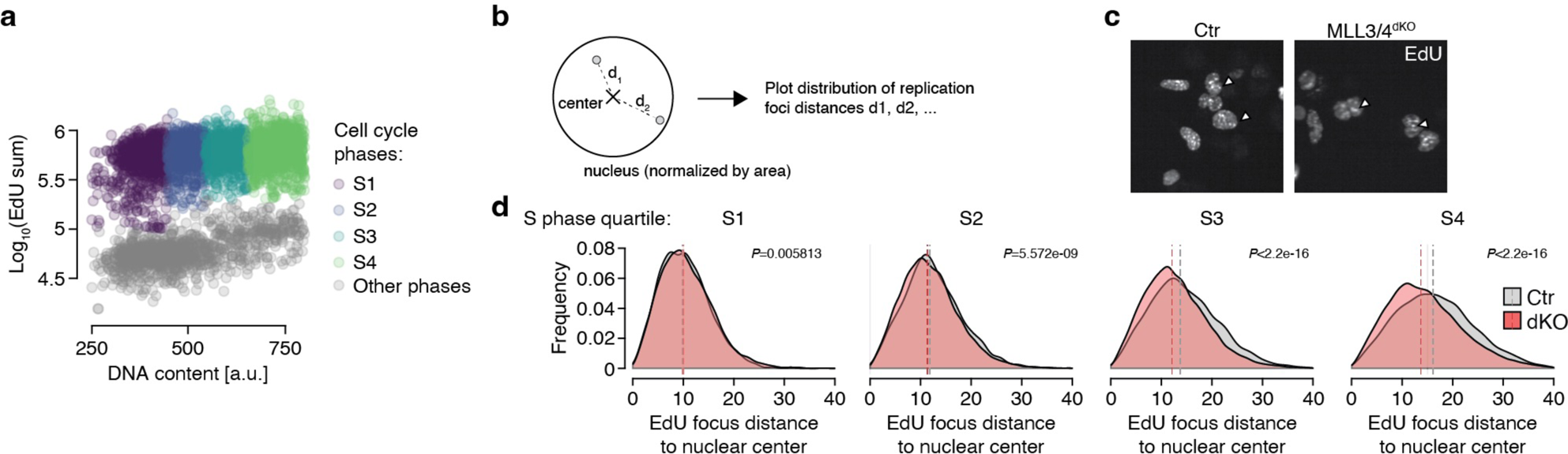
MLL3/4 impacts replication foci distribution. **a,** Scatter plot of EdU signal *vs* DNA content (based on DAPI staining) per MLL3/4^dKO^ or control ES cell derived from automated high content confocal microscopy. Colors indicate quartiles S1 to S4 of EdU^+^ cells based on DNA content. **b**, Scheme of measurement of replication foci distance to nuclear center. **c,** Example images of EdU foci from controls (MLL3^KO^) and MLL3/4^dKO^ cells at 20x magnification derived from high-content confocal microscopy. **d,** Histograms showing the distribution of replication foci distance to nuclear center in different S phase stages (quartiles based on DNA content; see a) in MLL3/4^dKO^ and controls (Ctr; MLL3^KO^). *P* values obtained through Wilcoxon Rank-Sum Test.

**Extended Data Figure 5:**
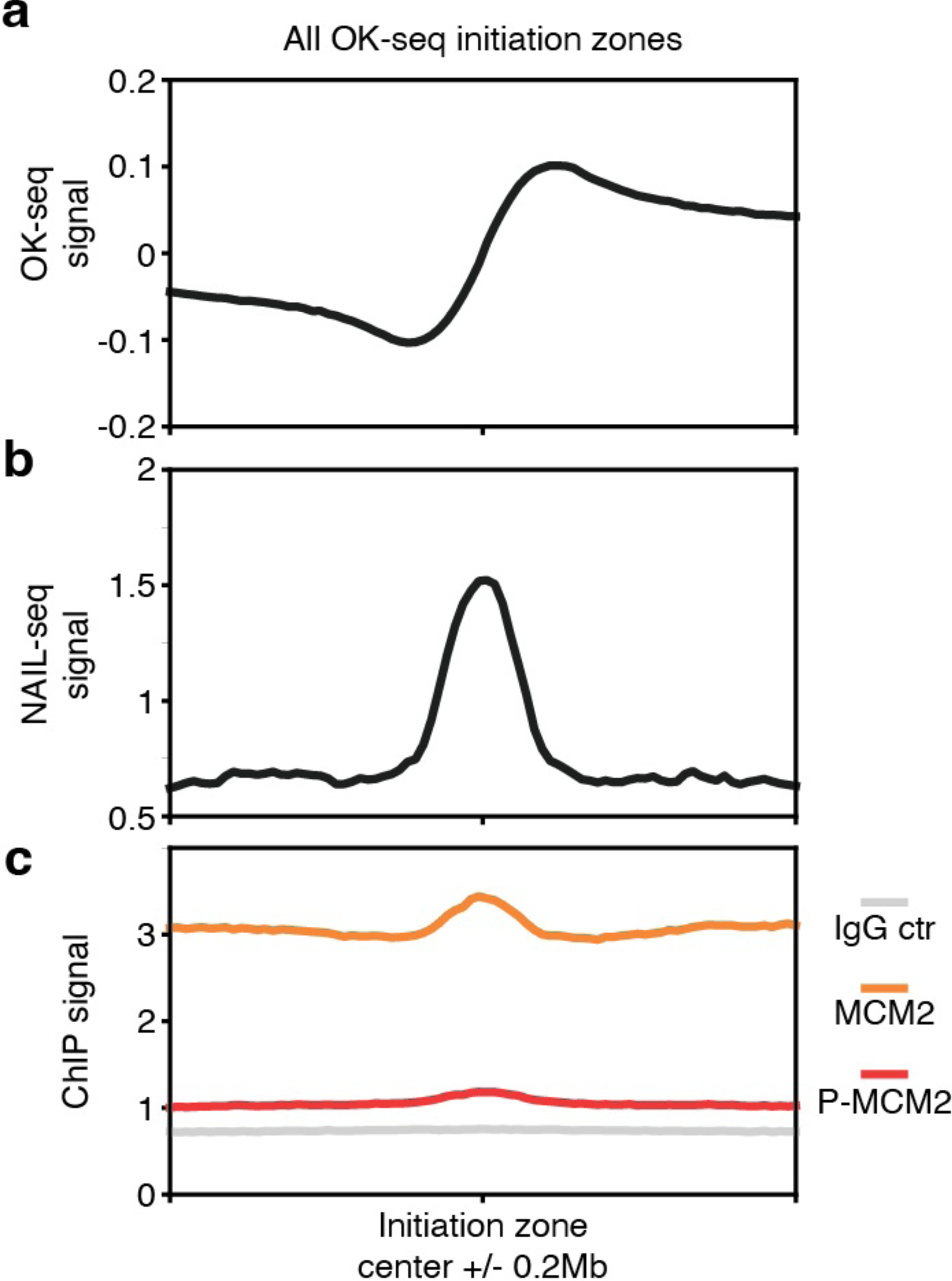
Validation of MCM2 ChIP-seq data. **a**-**c**, Metagene plots of OK-seq (a), NAIL-seq (b) and MCM2 (orange), phosphorylated MCM2 (red) and IgG control (gray) ChIP-seq data (c) centered at OK-seq initiation zones.

## EXTENDED DATA TABLES

**Extended Data Table 1:** Datasets used in this study.

| Type | Reference | Accession |
| --- | --- | --- |
| H3K4me1 C&T | This study | GSE216475 |
| H3K4me2 C&T | This study | GSE216475 |
| H3K4me3 C&T | This study | GSE216475 |
| H3K27me3 C&T | This study | GSE216475 |
| H3K9me2 C&T | This study | GSE216475 |
| H3K9me3 C&T | This study | GSE216475 |
| H4K20me1 C&T | This study | GSE216475 |
| H3K36me1 C&T | This study | GSE216475 |
| H3K36me2 C&T | This study | GSE216475 |
| H2BK5ac C&T | This study | GSE216475 |
| H3K9ac C&T | This study | GSE216475 |
| H3K14ac C&T | This study | GSE216475 |
| H3K27ac C&T | This study | GSE216475 |
| H4K8ac C&T | This study | GSE216475 |
| H4K16ac C&T | This study | GSE216475 |
| P-Pol II C&T | This study | GSE216475 |
| H2Az C&T | This study | GSE216475 |
| H3.3 C&T | This study | GSE216475 |
| γH2AX C&T | This study | GSE216475 |
| RAD21 C&T | This study | GSE216475 |
| H2BK119Ub C&T | This study | GSE216475 |
| H3K4me1 C&R | Boileau et al 2023 | GSE212950 |
| ATAC-seq | Chen et al 2018 | GSE93147 |
| NANOG ChIP-seq | Galonska et al 2015 | GSE56312 |
| H3K4me1 ChIP-seq | Chen et al 2018 | GSE93147 |
| OCT4 ChIP-seq | Buecker et al 2014 | GSE56138 |
| Pol II ChIP-seq | Marks et al 2012 | GSE23943 |
| MLL3/4 ChIP-seq | Dorigi et al 2017 | GSE98063 |
| H3K27ac ChIP-seq | Chen et al 2018 | GSE93147 |
| P300 ChIP-seq | Buecker et al 2014 | GSE56138 |
| SOX2 ChIP-seq | Galonska et al 2015 | GSE56312 |
| MCM2 ChIP-seq | This study | GSE216475 |
| P-MCM2 ChIP-seq | This study | GSE216475 |
| OK-seq | Petryk et al 2018 | GSE117274 |
| NAIL-seq | Liu et al 2021 | GSE174680 |
| BrdU-Repli-seq | Miura et al 2019 | GSE113985 |
| Bio-Repli-seq | This study | GSE216475 |
| RNA-seq | Boileau et al 2023 | GSE212950 |

